# Synthetic torpor in the rat protects the heart from ischaemia-reperfusion injury

**DOI:** 10.1101/2025.03.12.642790

**Authors:** Megan Elley, Ludovico Taddei, Muzammil Ali Khan, Una R Wilcox, Timna Hitrec, Martin Lewis, Anthony Pickering, Michael Ambler

## Abstract

During times of environmental stress, many animals enter torpor: a reversible protective physiological state typically characterised by reductions in core temperature, heart rate and oxygen consumption. Species that naturally enter this hypothermic and hypometabolic state are tolerant of ischaemia-reperfusion injury. Consequently, there is a growing interest in utilizing aspects of torpor for clinical applications, such as protection from stroke or myocardial infarction. It is currently unknown, however, whether a torpor-like state is protective in animals that do not naturally enter torpor. Using viral vector-mediated chemogenetic activation of the medial preoptic area of the hypothalamus, we induced synthetic torpor in the rat, a species that does not naturally enter torpor. We demonstrate this state is cardioprotective in an ex vivo ischaemia-reperfusion injury model with an ∼40% reduction in infarct size. Synthetic torpor-induced cardioprotection of the normothermic, isolated heart is not dependent on prior hypothermia in vivo. Phosphoproteomic analysis of cardiac tissue indicates the protective effects of synthetic torpor may be mediated by parallel activation of cell survival and stress tolerance pathways and inhibition of cell death pathways. These findings provide important insights into the mechanisms of organ protective effects of synthetic torpor states with implications for future clinical translation in humans.

## Introduction

Cardiovascular disease, including stroke and myocardial infarction, is the leading cause of death and disability globally^1,2^. Most current treatment strategies focus on re-establishing tissue perfusion, but there is interest in developing alternative strategies to improve tolerance to the ischaemia and/or reduce the impact of the reperfusion injury^3^. The idea that therapeutic hypothermia might reduce cell death during ischaemia-reperfusion has remained popular for many years, fuelled by encouraging animal studies^4–8^. However, despite early positive trials^9^, more recent clinical data do not support the hypothesis that induced hypothermia improves outcomes following cardiac arrrest^10^. Failure to translate benefits from animal studies to adult clinical conditions may be due to side-effects arising from the counter-regulatory reflexes triggered by physical cooling. These reflexes lead to haemodynamic instability, shivering with increased oxygen consumption, cardiac arrhythmias, and electrolyte and glucose disturbances^11,12^. Hence, interest is growing in centrally driven hypometabolic, hypothermic states that might avoid these counter-regulatory responses. With this in mind, researchers have turned to hibernating animals as a model for a centrally driven protective hypometabolic state^8,13–15^.

During hibernation, animals enter torpor, which is a remarkable physiological adaptation typically characterised by rapid and profound reductions in core temperature, heart rate, and oxygen consumption^16^. It serves as a protective adaptive response to relative energy deficit, achieved by a controlled reduction in metabolic demand in response to reduced availability of substrate. Torpor can be prolonged, in seasonal hibernators, or brief in daily heterotherms such as the mouse. During torpor, body temperature can be lowered to a few degrees above ambient temperature^17,18^. Metabolic activity may fall to below 5% of the euthermic rate, with active and profound decreases in cardiac output, and minute ventilation, all of which rapidly return to ‘normal’ values on arousal without harm to the individual^17,18^. Seasonal hibernators spend prolonged periods of time torpid and are highly tolerant of ischaemia-reperfusion injury^13,19,20^. This has motivated efforts to develop biomimetic torpor-like states for clinical applications that could provide a range of benefits including protection from ischaemia-reperfusion injury^15^.

A fundamental hypothesis underlying interest in such translational torpor research is that conserved mechanisms exist in species that do not naturally enter torpor (potentially including humans), aspects of which might be harnessed to improve clinical outcomes. There are several reasons to suppose this to be the case. Firstly, torpor is extremely widespread, seen in all endothermic vertebrate classes, including primates^18,21^. Secondly, a unique torpor gene (or complement of genes) has so far not been described. Rather, those mammals that enter torpor do so through activation of ubiquitous genetic pathways^22–24^. Thirdly, body temperature is not fixed even in ‘strict’ homeotherms: non-rapid-eye-movement sleep (NREM) is a controlled state of reduced body temperature and inactivity, features it shares with torpor^25,26^.

Attempts to induce a “synthetic” torpor-like state in a species for which it is not a natural behaviour have focused on the rat, and include inhibition of the rostral ventromedial medulla^27^; peripheral infusion of drugs, such as adenosine and neurotensin analogues^28,29^; and targeted activation of neurons in the preoptic area of the hypothalamus^30^. Of these approaches, the latter is arguably the closest to mimicking natural torpor, since it engages neurons in part of the hypothalamus that drives entry into natural torpor in mice^30–32^. We sought to characterise the state of synthetic torpor induced in the rat by targeted activation of neurons in the hypothalamic medial preoptic area (MPA), and to test the hypothesis that induction of synthetic torpor protects the heart from ischaemia-reperfusion injury.

## Results

### Induction of biomimetic synthetic torpor in the rat

We used viral vector-mediated chemogenetics to selectively activate neurons in the MPA (figure 1A). The vector (AAV-CaMKIIa-hM3D(Gq)-mCherry) was injected bilaterally into the rat MPA, resulting in local expression of the modified receptor hM3Dq (a Gq-coupled Designer Receptor Exclusively Activated by Designer Drug, DREADD^33,34^) under the control of the CaMKIIα promoter^35,36^ (producing MPA^Gq^ rats). Control animals underwent identical procedures but received injection of AAV-CaMKIIa-EGFP into the MPA which results only in the expression of a green fluorescent protein (MPA^EGFP^ rats).

**Figure 1.**
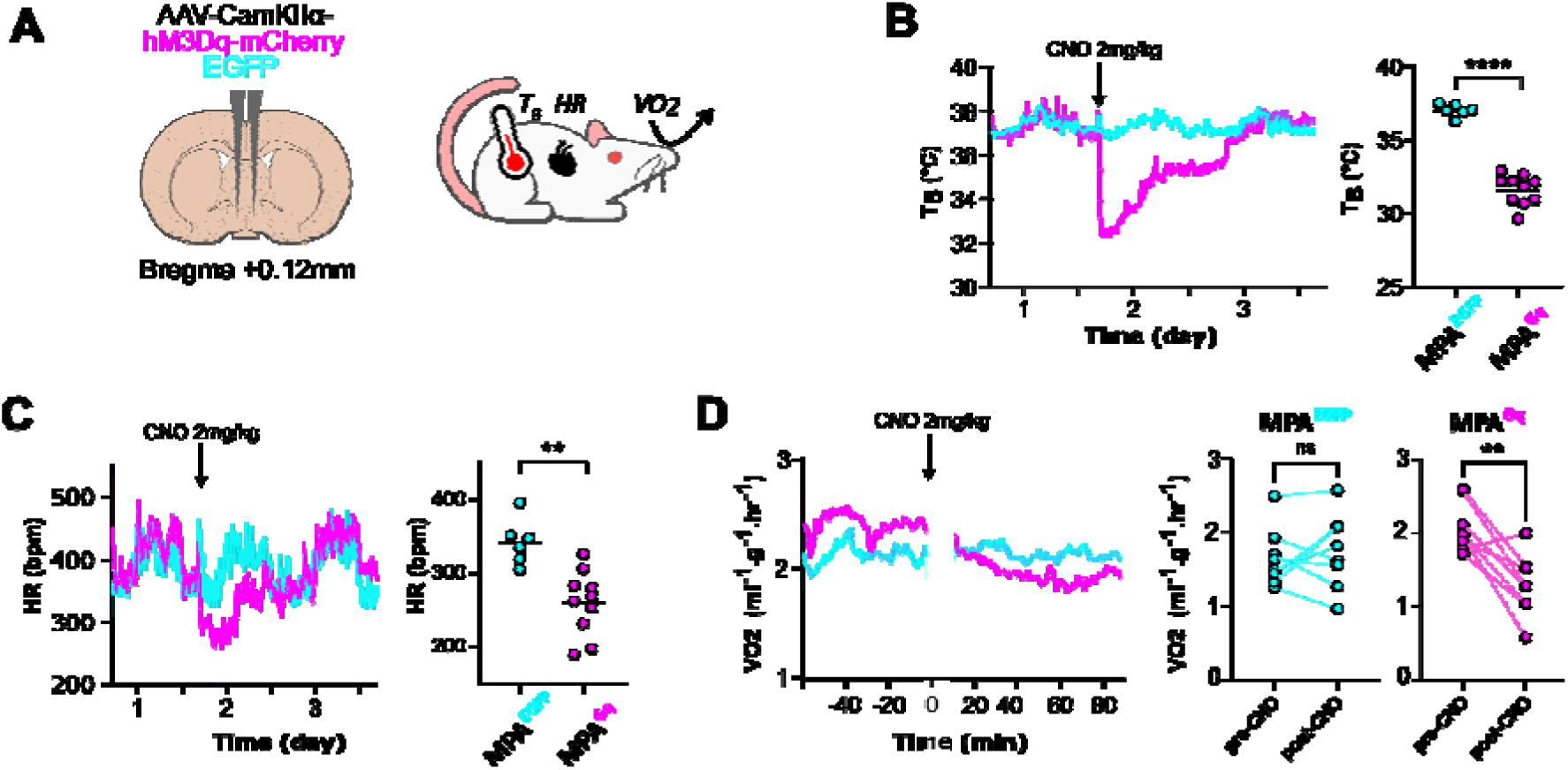
Chemogenetic activation of neurons in the rat medial preoptic area (MPA, A) induces synthetic torpor. characterised by reduced core body temperature (B, F(3,24) = 106.4. *p* < 0.0001), bradycardia (C, F(3,24) = 9.637, *p* = 0.0002), and reduced oxygen consumption (D, MPA^GFP^ t(7) = 0.5655, *P* = 0.5894, MPA^Gq^ t(7) = 4.305, *P* = 0.0035), For B and C, *n* = 6 controls and 10 synthetic torpor, mean value for 90-100 minutes after CNO 2mg/kg IP. For D, *n* = 8 MPA^GFP^ and 8 MPA^Gq^ pre (50-60 minutes) vs post (120-130 minutes) CNO. Shaded area indicates S.E.M. * indicates *p* <0.05, ** indicates *p* < 0.01, *** indicates *p* < 0.001, **** indicates *p* < 0.0001. Abbreviations: T_B_, core temperature; VO_2_, oxygen consumption; HR, heart rate; CNO, clozapine-N-oxide; ST, synthetic torpor; bpm, beats per minute.

Subsequent MPA neuronal activation using intraperitoneal CNO (2 mg/kg) induced a rapid and reversible decrease in core temperature (mean temperature post-CNO in MPA^Gq^ rats 31.64 ± 1.02°C vs MPA^EGFP^ rats 37.05 ± 0.43°C, p <0.0001, see Figure 1B). This was accompanied by a 24% lower heart rate (mean heart rate post-CNO in MPA^Gq^ rats 260 ± 44 bpm vs MPA^EGFP^ 342 ± 32 bpm, p <0.005, figure 1C). This was also associated with a 39% reduction in oxygen consumption in MPA^Gq^ rats (mean oxygen consumption prior to CNO 2.08±0.34ml/g/hr vs 1.29±0.42ml/kg/hr post-CNO, p =0.0035, figure 1D). Whereas there was no significant change in oxygen consumption in MPA^EGFP^ rats (mean oxygen consumption prior to CNO 1.25±0.4ml/g/hr vs 1.68±0.62ml/g/hr post-CNO, p =0.589, figure 1D). Hence, chemoactivation of neurons in the rat MPA produces a synthetic torpor state that recapitulates three key features of natural torpor: hypothermia, bradycardia, and reduced oxygen consumption.

### Mapping the distribution of the neurons responsible for synthetic torpor in the rat

We mapped the location of mCherry-expressing transduced neurons within the anterior hypothalamus of 6 animals (supplementary figure 2). We observed transduced neurons in a region stretching from bregma +0.12mm to -0.48mm, spreading 1mm bilaterally from the midline, and from the ventral surface of the hypothalamus to 1.5mm dorsally (Figure 2F). This region includes the targeted MPA as well as more caudal regions of the preoptic area such as the medial preoptic nucleus.

**Figure 2.**
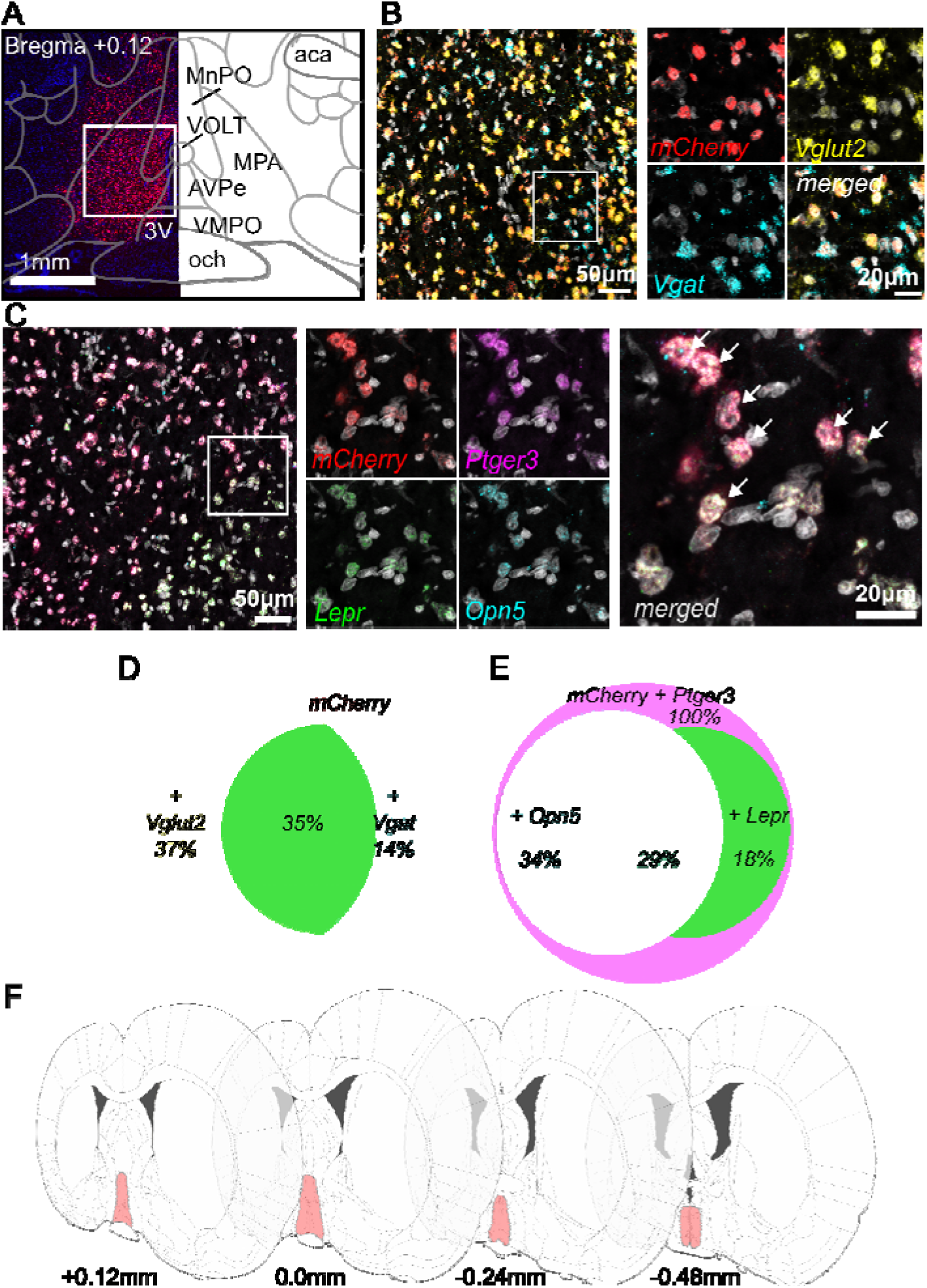
Characterisation of neurons in the rat MPA responsible for inducing synthetic torpor. A, viral vector transduction with mCherry in the MPA (red) and corresponding anatomical schematic. B, RNA *in-situ* hybridisation labelling vector transduced cells (*mCherry*, red), and those expressing *Vglut2* (yellow) and/or *Vgat* (turquoise). C, RNA *in-situ* hybridisation labelling vector transduced cells (*mCherry*, red), and those expressing *Ptger3* (purple), *Lepr* (green), and *Opn5* (turquoise). White arrows indicate transduced cells that express *Ptger3*, *Lepr*, and *Opn5*. D, co-expression of *Vglut2* and *Vgat* amongst transduced cells. E, expression of QPLOT markers amongst vector transduced cells (note all transduced cells expressed *Ptger3*). F, schematic showing extent of viral vector transduction common to all animals. Abbreviations: MPA, medial preoptic area; MnPO, median preoptic nucleus; VOLT, vascular organ of the lamina terminalis; 3V, 3^rd^ventricle; AVPe, anteroventral periventricular nucleus; VMPO, ventromedial preoptic nucleus; och, optic chiasm; aca, anterior commissure (anterior part); *Vglut2*, vesicular glutamate transporter 2; *Vgat*, vesicular GABA transporter; *Ptger3*,

### Characterising the neurons in the rat MPA responsible for inducing synthetic torpor

To establish whether the neurons involved in inducing this synthetic torpor state are excitatory or inhibitory, we used in situ hybridization (ISH) to examine colocalization of vector-mediated mCherry expression (figure 2A-E) with genes that encode vesicular glutamate transporter 2 (Vglut2, gene ID Sl17a6) and vesicular GABA transporter (Vgat, gene ID Slc32a1) (Figure 2B). We found a large proportion of transduced neurons expressed Vglut2 (72%, Figure 2D). We also detected a population of transduced neurons that co-express both Vglut2 and Vgat (35%), termed ‘dual-phenotype’ neurons^37^ (Figure 2D). A small proportion of vector transduced neurons were inhibitory (14%), expressing Vgat and a remainder expressed neither Vglut2 nor Vgat (15%) (figure 2D).

Previous studies have identified a population of neurons within the preoptic area of the mouse hypothalamus, which can integrate stimuli to regulate thermogenesis and metabolism and induce a torpor-like state upon activation^38^. These stimuli include temperature changes both locally within the hypothalamus and from peripheral signals in the skin, inflammatory prostaglandin signalling, leptin, oestrogen, and violet light. These neurons were called QPLOT neurons, reflecting their co-expression of neuropeptide pyroglutamylated RFamide peptide (Qrfp), Prostaglandin EP3 receptor (encoded by Ptger3), Leptin receptor (encoded by Lepr), violet light-sensitive opsin, Neuropsin (encoded by Opn5) and neurokinin B (encoded by Tacr3). Consequently, we sought to determine whether the neurons responsible for inducing synthetic torpor in the rat share similarities to this torpor-inducing population identified in mice. We again used ISH to assess colocalization of vector-expressed mCherry with three markers of QPLOT neurons: Ptger3, Lepr, and Opn5. We found 100% of the virally transduced cells also expressed Ptger3 (figure 2D-E). Additionally, 34% of these mCherry & Ptger3 positive neurons also expressed Opn5, 18% co-expressed LepR (figure 2E), and 29% of mCherry positive neurons were positive for all three of the five QPLOT neuron markers we labelled (figure 2E). These findings suggest a degree of conservation of neuronal organisation within the MPA between mice and rats and are consistent with synthetic torpor being produced by activation of analogous “torpor-generating” circuits across species.

### Induction of synthetic torpor protects the heart from ischaemia-reperfusion injury

Both hypothermia^39^ and torpor^13,19,20^ have been shown to protect against ischaemia-reperfusion injury. We therefore sought to test the effects of synthetic torpor on tolerance to cardiac ischaemia-reperfusion injury in the ex vivo Langendorff isolated heart preparation^40^ (Figure 3A). Induction of synthetic torpor, 90 minutes prior to removal of the heart for the Langendorff preparation and exposure to ischaemia-reperfusion, reduced infarct size by 39.5% (percentage infarcted myocardium: MPA^Gq^ 18.7 ± 9.6% vs MPA^EGFP^ 30.9 ± 9.1%, p = 0.039, Figure 3B & C). Importantly, although the synthetic torpor state reduced the body temperature of the rats prior heart excision, the perfusion of warm Krebs-Henseleit Buffer (KHB) to the cardiac tissue for a 30-minute stabilisation period ensured that there were no differences in temperature at the time of ischaemia and reperfusion.

**Figure 3.**
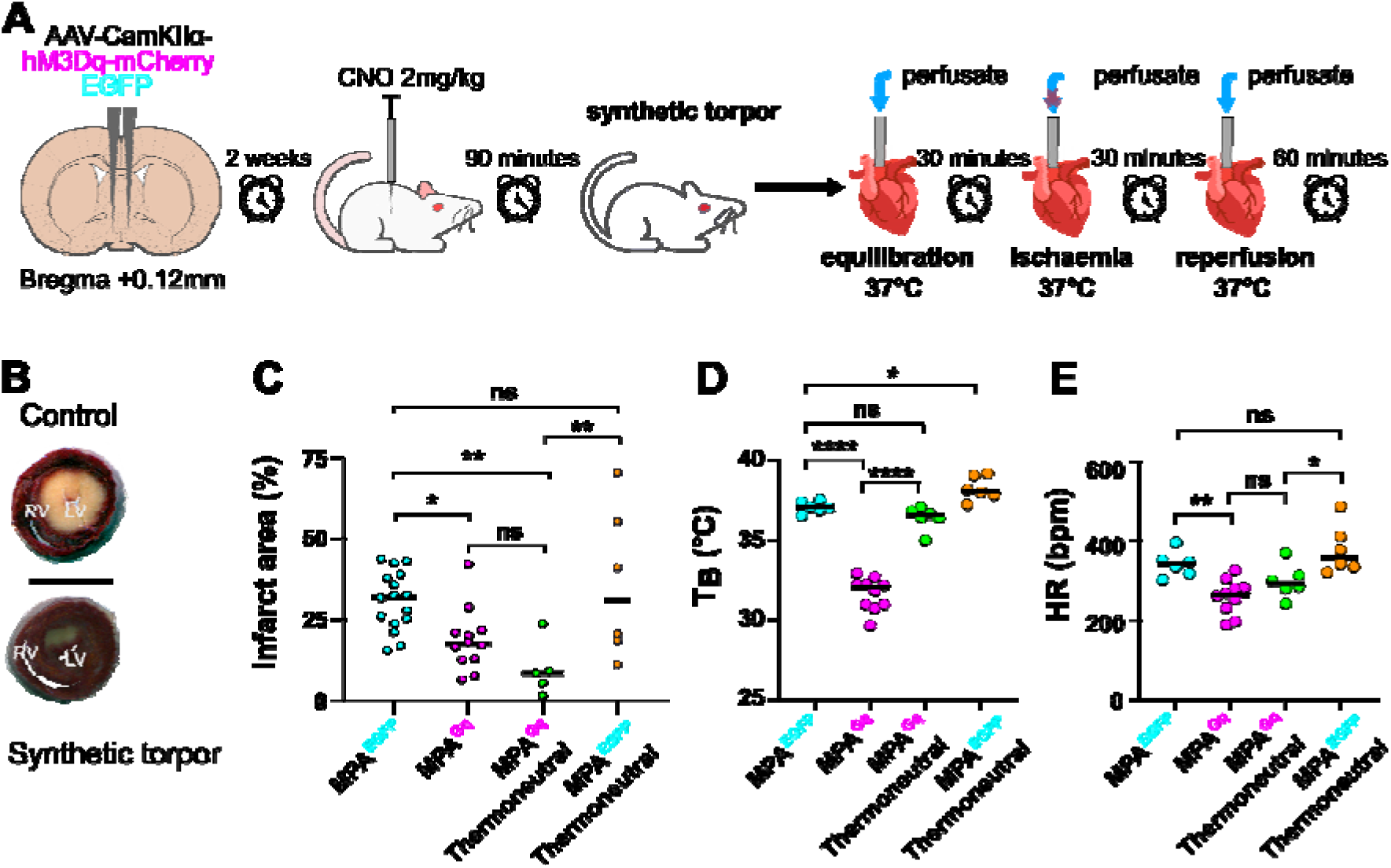
Synthetic torpor protects the heart from *ex vivo* ischaemia-reperfusion injury. Rats first underwent bilateral injection of vector (hM3DGq or GFP) into the MPA. After two weeks, they all received CNO, inducing synthetic torpor in rats that had received hM3DGq vector, followed by removal of the heart and ischaemia-reperfusion in the Langendorff preparation (A). Pre-conditioning with synthetic torpor resulted in significantly reduced infarct size as measured by 2,3,5-Triphenyltetrazolium Chloride (TTC) staining. (B, C). Scale bar represents 1cm. For C, Controls (n = 16), torpor (n = 12), thermoneutral synthetic torpor (n = 5), & thermoneutral controls (n = 6). This cardioprotective effect was not dependent on a reduction in core temperature as rats held at a thermoneutral ambient temperature following CNO injection were still protected (C, ANOVA F (3,35) = 6.542, *p* = 0.0013) MPA^EGFP^ compared to MPA^Gq^ *p* = 0.039 and thermoneutral MPA^EGFP^ compared to thermoneutral MPA^Gq^ *p* = 0.007. Inducing synthetic torpor in a thermoneutral environment prevented the drop in core temperature (D, F(3,24) = 106.4. *P* < 0.0001), but not the bradycardia (E, F(3,24) = 9.637, *p* = 0.0002). Holm-Sidak correction for multiple comparisons. Data shown is median and IQR. For D & E, MPA^EGFP^ (n = 6), MPA^Gq^ (n = 10), MPA^Gq^ thermoneutral (n =6) and MPA^EGFP^ thermoneutral (n = 6), mean value for 90-100 minutes after CNO 2mg/kg IP * = *p* <0.05, ** = *p* < 0.01, **** = *p* < 0.00001, ns = *p* >0.05. Abbreviations: CNO, clozapine-N-oxide; RV, right ventricle; LV, left ventricle; T_B_, core temperature; HR, heart rate; bpm, beats per minute.

To examine whether the synthetic torpor-induced myocardial protection depends upon the preceding hypothermia, we housed groups of MPA^Gq^ and MPA^EGFP^ rats at a thermoneutral temperature of 28-30°C. Administration of CNO to MPA^Gq^ rats under thermoneutral conditions prevented the drop in core temperature that was observed in recordings at room temperature (p <0.0001, see Figure 3D and Supplementary Figure 1A). Furthermore, comparing thermoneutral MPA^Gq^ rats with room temperature MPA^EGFP^ control rats showed no significant temperature change (mean temperature post-CNO in thermoneutral MPA^Gq^ rats 36.4 ± 0.73°C vs MPA^EGFP^ 37.05 ± 0.43°C, p =0.44, Figure 3D). However, the bradycardia observed with synthetic torpor induced at room temperature persisted even when rats were housed at a thermoneutral temperature (Figure 3E). A 21% lower heart rate was observed in the thermoneutral MPA^Gq^ compared to MPA^EGFP^ rats (mean heart rate post-CNO in thermoneutral MPA^Gq^ rats 298 ± 43 bpm vs thermoneutral MPA^EGFP^ rats 379 ± 62 bpm, p <0.05, see figure 3E and supplementary figure 1B). This indicates that the heart rate changes associated with synthetic torpor were not secondary to the induced hypothermia. Additionally, there were no significant differences in the heart rate of animals in the MPA^EGFP^ condition compared to the thermoneutral MPA^EGFP^ condition (p = 0.53, Figure 3E) or between the MPA^Gq^ and thermoneutral MPA^Gq^ condition (p = 0.38, figure 3E and supplementary figure 1B).

We found that thermoneutral synthetic torpor still reduced the infarct size (percentage infarct in the thermoneutral MPA^Gq^ rats 9.6 ± 8.4 % compared to thermoneutral MPA^EGFP^ rats 36.2 ± 23.5 %, p = 0.005and room temperature MPA^EGFP^ rats p = 0.007, Figure 3C). There were no significant differences in infarct size between the MPA^Gq^ and thermoneutral MPA^Gq^ groups, (p = 0.31) nor between the MPA^EGFP^ and thermoneutral MPA^EGFP^ groups, (p = 0.38) (figure 3C). We also found no correlation between individual nadir temperature during synthetic torpor and infarct size (p = 0.3714, R^2^ = 0.089, Supplementary figure 5). These findings indicate that the myocardial protection conferred by synthetic torpor is independent of the hypothermia.

Heart rate was monitored during the Langendorff assay to test for the possibility that the bradycardia observed in vivo during synthetic torpor persisted in the ex vivo cardiac preparation and hence contributed to the myocardial protection. No differences were observed in hearts rates during the 30-minute equilibration period (prior to induction of ischaemia) in MPA^Gq^ (240 ± 28 bpm), MPA^EGFP^ (248 ± 23 bpm), thermoneutral MPA^Gq^ (225 ± 20 bpm) and thermoneutral MPA^EGFP^ (225 ± 21 bpm) conditions, F(3,19) = 1.35, p = 0.29 (Supplementary Figure 1C). This indicates the decreased heart rate observed during synthetic torpor in vivo does not persist ex vivo.

### Phosphoproteomic analysis of cardiac tissue following synthetic torpor

To further understand the mechanisms underlying the cardioprotective effect observed in synthetic torpor, we conducted mass spectrometry (MS) using tandem mass tag (TMT) which allowed us to analyse the phosphoproteomic profile of rat heart tissue following 90 minutes of synthetic torpor (without ischaemia-reperfusion injury exposure) compared to control rats. After quality control and preprocessing, we identified 4064 phosphopeptides from 1090 unique genes in our rat samples (supplementary figure 3). Missingness, inter-sample correlation, PCA distance, and phosphosite distributional properties did not identify any outlier samples (supplementary figure 4). The purpose of this analysis was exploration and hypothesis generation and therefore the results were not corrected for multiple comparisons (figure 4).

**Figure 4.**
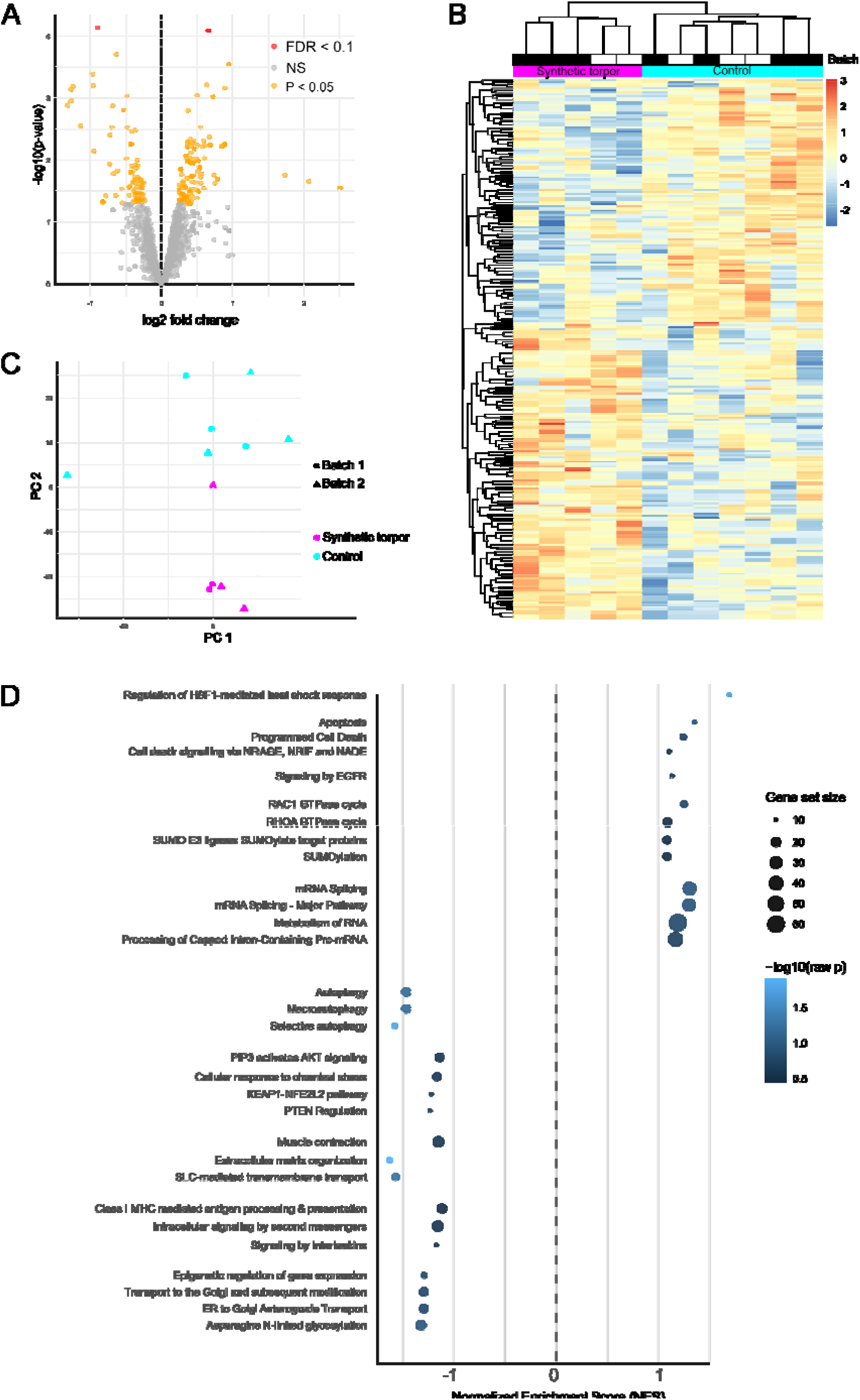
Phosphoproteome analysis of the heart identifies gene pathways associated with synthetic torpor. ,Volcano plot showing differentially expressed phosphopeptides after limma modelling (A). Heatmap shows the top 300 differentially expressed phosphosites between synthetic torpor (n = 5) and controls (n = 7) after limma modelling,rows (phosphosites) and columns (sample) clustered by similarity of expression so that phosphopeptides that covary cluster and samples that covary also cluster toether.(B). Principal component analysis confirms group separation and effective batch correction (C). Gene-level set enrichment analysis (GSEA) performed using a single representative phosphosite per gene (selected based on limma t statistic) identifies key pathways modulated by entry into synthetic torpor (D).

We analysed the proteomics data in two parallel and complementary ways. One approach collapsed the phosphosite data from multiple potential sites to a single site per protein, with representative sites selected by limma^41^ t-statistic for the most differentially enriched between synthetic torpor and controls. The corresponding gene for each protein (1073) and the associated t-statistic was then used as the input for Gene Set Enrichment Analysis (GSEA) using ReactomePA^42^,which returned 136 differentially enriched pathways. GSEA performed in this way on phosphoproteome data is agnostic to the direction of modulation (that is, while a negative enrichment score indicates relative dephosphorylation of the representative phosphosite, GSEA does not impute the effect this will have on the pathway in terms of increasing or decreasing activity). We focused on the top 30 most differentially enriched pathways. This analysis revealed a coordinated ensemble of pathway modulation involving control of apoptosis and autophagy, cellular stress response mechanisms, control of transcription and translation, and regulation of heat shock proteins (figure 4).

The second approach reconstructed rat phosphoprotein structure from phosphopeptides, mapped them to human orthologues via their corresponding gene^43,44^, and then performed post-translational modification set enrichment analysis (PTM-SEA). This approach, by retaining site specific phosphorylation state data, identifies key kinases by the signature pattern of phosphorylation changes across the analysed proteome, in this case returning whether the kinase has been activated or inhibited. We were able to fully map 1384 phosphosites from 595 proteins from our rat sample to corresponding human phosphoproteins. This is necessary because the rat phosphoproteome is less well studied and the PTM-SEA package is based on human data^45^. We found 156 differentially enriched kinase signatures across the 1384 phosphosites. We focused on the 15 most differentially enriched kinase signatures (figure 5). Again, we saw a coordinated ensemble of kinase modulation by synthetic torpor, including: suppression of mTOR-Akt-ribosomal growth signalling indicating reduced anabolic drive (reduced Akt, p70S6K, RPS6KA5); dampened stress-activated MAPK pathways (MAPK12-14 and MAPKAPK2,3, & 5); downregulated excitability (reduced activity of MAPK10, CAMK2A & CAMK2G); reduction of cellular energy consumption (reduced AMPKα1); and reduced growth/hypertrophy (reduced ACVR1 activity). In parallel we saw increased activity of CSNK2A2, suggesting increased stress tolerance and cytoprotection. Hence, both gene-level pathway analysis and specific kinase enrichment analysis indicate that synthetic torpor invokes a coordinated pattern of protective adaptations.

**Figure 5.**
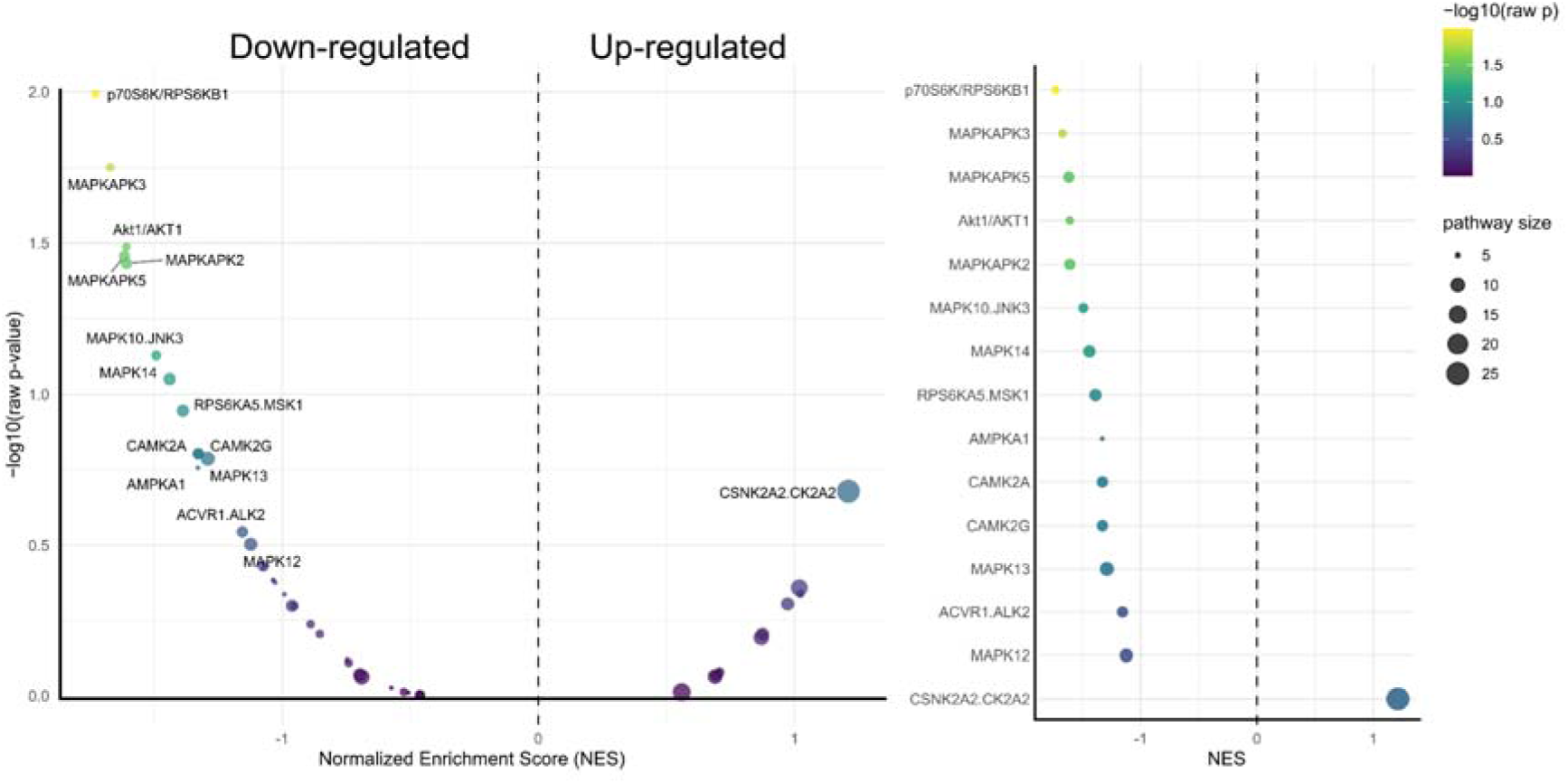
Site-level phosphoproteome analysis. After mapping from rat to human phosphoproteins, post-translational set enrichment analysis (PTM-SEA) was performed on 1384 aligned phosphosites from 595 rat gene-human protein pairs. Kinase signatures are shown ranked by normalized enrichment score (NES); the top 15 by absolute NES are labelled for clarity. Colour represents −log10(raw p-value), and point size indicates pathway size.

## Discussion

We implemented a chemogenetic model of synthetic torpor in the rat^46^, a species that does not naturally enter torpor, by targeting a population of hypothalamic neurons analogous to those that drive fasting-induced torpor in the mouse. We demonstrate that synthetic torpor in the rat is strongly cardioprotective in an ex vivo ischaemia–reperfusion injury model, even in the absence of hypothermia. In parallel, phosphoproteome analysis revealed changes consistent with enhanced cellular stress tolerance and inhibition of cell death pathways, which may contribute to the observed cardioprotection. Importantly, because synthetic torpor was induced in a species that does not naturally enter torpor, these findings support the existence of conserved, cross-species mechanisms that could potentially be translated for therapeutic use in humans.

Recent years has seen a resurgence of interest in torpor and torpor-like states, driven by a hope that aspects of this remarkable physiology could be mimicked in both clinical and space-flight situations. Here we demonstrate that chemogenetic activation of neurons in the MPA of a non-torpid species, the rat, recapitulates three main features of natural torpor: hypothermia (Figure 1B), bradycardia (Figure 1C) and reduced oxygen consumption (Figure 1D). Interestingly, we did not observe a significant reduction in the motor activity of the rats during synthetic torpor (Supplementary Figure 6) which is typically observed in mice during both natural and synthetic torpor ^31^. The preoptic area of the hypothalamus (POA) is a key centre for control of energy balance, body temperature, and as a driver of natural fasting-induced torpor in the mouse^31,46–48^. Analysis of pooled single-nucleus RNA sequencing data has attempted to phenotype the critical neuronal population, from which emerge ‘QPLOT neurons’^31,38,49^. These cells express QRFP^46^, prostaglandins E3 receptor (EP3)^50^, leptin receptor^51^, the violet light sensitive receptor Opn5^52^, and neurokinin 3 receptors (coded for by Tacr3)^53^. Expression of two or more of these markers is considered sufficient to identify neurons from this population^54^. 81% of the neurons we targeted with the DREADD viral vector expressed either two or three of these QPLOT neuron markers (figure 2C & E). And 29% of these neurons expressed all three markers used, which included Ptger3, Lepr and Opn5 (figure 2C & E). The observation that virally transduced cells in the rat MPA also express QPLOT markers indicates the presence of a neuronal population in rats with expression patterns and physiological effects equivalent to those described in mice, supporting the existence of a conserved, cross-species population.

We found that 72% of transduced mCherry neurons were positive for Vglut2, a marker of excitatory neurons (figure 2B & D). Of these neurons, 35% were labelled as ‘dual-phenotype’^37^, expressing markers of both excitatory and inhibitory neurons (Vglut2 and Vgat, figure 2B & D). This was anticipated, as we used viral vector-mediated chemogenetics under the control of the CamKIIα promoter, which targets excitatory neurons^55,56^, but with some tropism for inhibitory neurons also^57^. Furthermore, a subset of neurons involved in mouse torpor, express both excitatory and inhibitory RNA markers^46^. Whether these neurons are functionally release glutamate and GABA remains to be seen, but evidence in the rat suggests that while EP3 receptor-expressing thermoregulatory neurons in the preoptic area may express excitatory and inhibitory markers, they are functionally inhibitory^58^

A key question that emerges from the observation of synthetic torpor in the rat is: what is the natural role of these neural circuits and why do they not naturally induce torpor in response to fasting? EP3 receptor expressing neurons in the rat POA have been implicated in bidirectional control of body temperature^58^. These neurons are inhibited by prostaglandins to produce fever and activated by warmth exposure to reduce body temperature. It is likely that these neurons and the population we have identified overlap. Hence, rats might maintain the capacity to activate these neurons in response to rising ambient temperature (and inhibit them in response to prostaglandins) but no longer be able to activate them in response to fasting. Alternatively, the rat may engage these neurons to reduce temperature at sleep onset^59^, but may have lost the ability to engage them to the extent that the animal enters torpor. Torpor and sleep may be functionally linked: mice enter and emerge from torpor through NREM sleep^60^ and the medial preoptic area of the hypothalamus appears to be responsible for triggering natural torpor^30,31,47^ as well as the cooling associated with NREM sleep onset^25,26^.

Animals that were pre-conditioned with 90 minutes of synthetic torpor showed a 40% smaller infarct size compared to control animals (figure 3). These findings support prior studies in species known to naturally enter torpor. For example: seasonal hibernators in torpor are highly tolerant of ischemia-reperfusion injury in multiple organs, including kidney and intestines^61–64^; inducing a torpor-like state using deep brain stimulation (DBS) to stimulate the medial preoptic area in mice is neuroprotective in a brain ischaemia model^7^; and targeting RFP-amide expressing neurons in the MPA to trigger a torpor-like state ameliorates acute kidney injury^65,66^ following ischaemia-reperfusion. Until now, this work was all carried out in species known to naturally enter torpor (mice and ground squirrels) and so it was not clear whether other species might also benefit from an induced torpor-like state. Our study demonstrates for the first time that inducing a torpor-like state in a species (the rat) that does not naturally enter torpor is protective in a cardiac ischemia-reperfusion injury model.

This study utilized an ex vivo model of ischaemia-reperfusion injury in the Langendorff preparation. Our finding that activation of MPA EP3-expressing neurons in the rat was cardioprotective indicates that activity in this neuronal population induces changes in the cardiomyocytes that persist when the heart is isolated from the rest of the organism. Whether the protective signal in vivo is circulating or released locally via vagal efferents remains an important question. The bradycardia we observed in vivo in response to chemoactivation of neurons within the MPA still occurs in the absence of hypothermia (figure 3E and supplementary figure 1B), suggesting active suppression of heart rate via the cardiac vagus nerve. Importantly, bradycardia did not persist during the ex vivo Langendorff preparation (supplementary figure 1C). This was important to assess, as lower heart rate in the Langendorff assay has previously been shown to decrease the size of the infarcted tissue area^67^. Hence, our model of synthetic torpor appears to engage protective responses that persist in isolation and are neither temperature-dependent, nor driven by a simple reduction in the heart rate during ischaemia-reperfusion.

Strikingly,chemogenetic activation of MPA neurons under thermoneutral conditions (which prevented hypothermia), still reduced infarct size. This was established by the thermoneutral condition, which prevented any decrease in core body temperature while the animals were in synthetic torpor (figure 3D) yet still resulted in a significant decrease in myocardial infarction size compared to control (figure 3C). We also observed no correlation between individual nadir surface temperature during synthetic torpor and the infarct size. These data show that the hypothermic component of torpor is not necessary for the protective effects of synthetic torpor. There have so far been mixed findings regarding the necessity for a drop in temperature to observe protection. One study used deep brain stimulation or chemogenetics to activate neurons within the mouse MPA to drive a torpor-like state and found that hypothermia is required for neuroprotection in a stroke model^65^. In contrast, and similar to our present study, Kyo and colleagues demonstrated that the hypothermic element of a torpor-like state in the mouse is not required to reduce acute kidney injury in response ischaemia^66^. Torpor-induced reduction in core temperature might be sufficient to improve tolerance to ischaemia-reperfusion injury, but some models (our present study and that of Kyo et al^66^) might engage additional, temperature-independent protective responses.

To explore the potential mechanisms occurring at the cardiac cellular level during synthetic torpor, we analysed the phosphoproteome of hearts taken from animals in synthetic torpor. We found many differentially regulated phosphoproteins and enrichment of numerous pathways associated with cell stress response, cell cycle and death (figure 6). The PTM-SEA data (figure 5) suggests activation of CSNK2A2 in response to synthetic torpor, a protein kinase associated with promoting cell survival, DNA repair, and inhibition of apoptosis^68^. A number of kinases were also inhibited, including Akt and p70S6K, which are associated with the mTOR signalling pathway^69^. mTOR signaling is implicated in metabolic regulation, and a decrease in activity of this pathway indicates reduced anabolism^69^, which is also observed in calorie restriction^70^ (a known natural driver of daily torpor^71^) and altered mTOR signaling is also seen in deep hibernating animals such as ground squirrels^72,73^. Inhibiting mTOR signaling protects against myocardial ischemia-reperfusion injury, by regulating autophagy and limiting cardiomyocyte apoptosis^74,75^.Decreased AMPK signaling is also indicative of a shift in cellular metabolism, playing an integral role in coordinating autophagy and mitochondrial function^76^.

**Figure 6.**
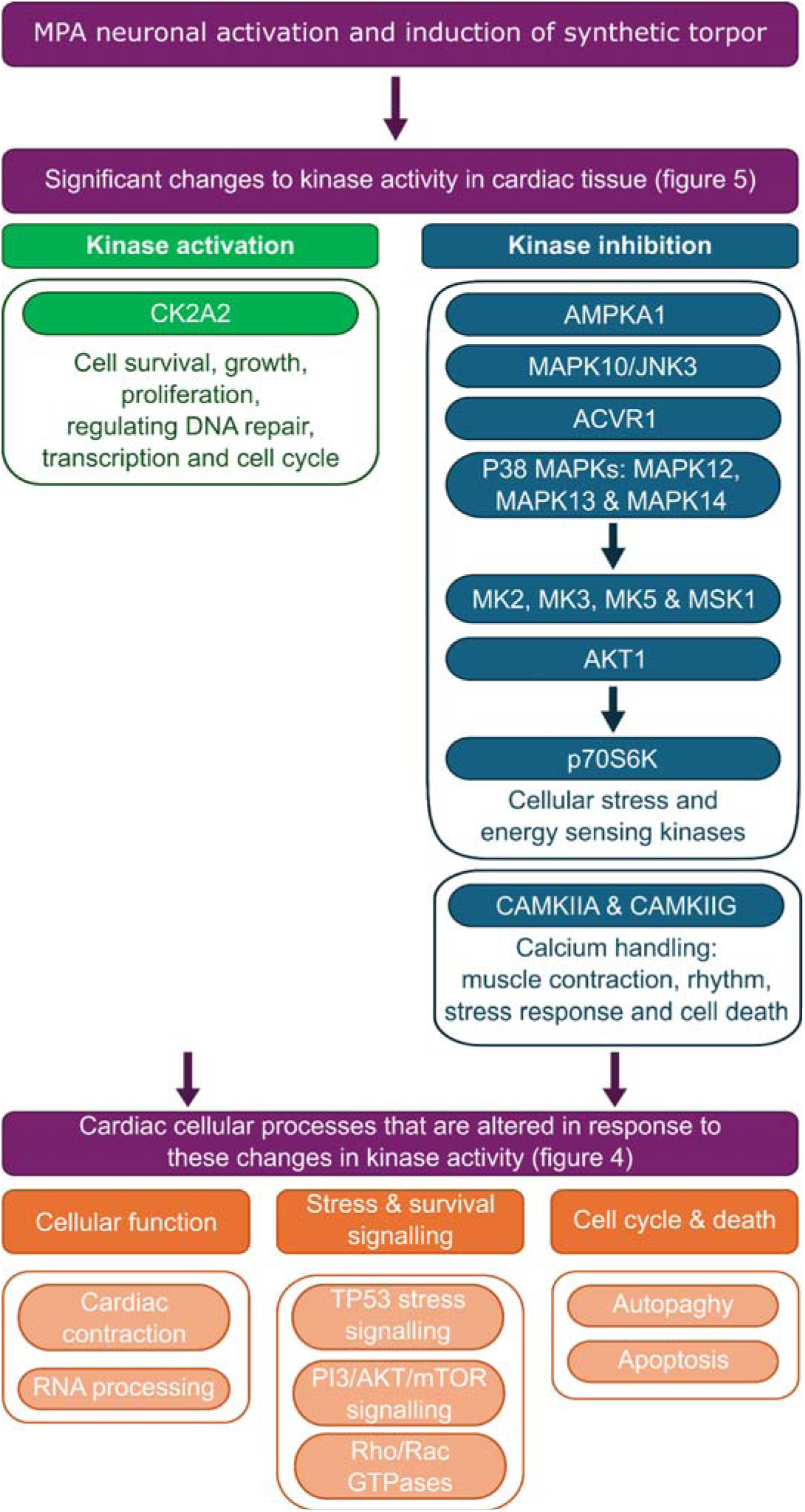
Changes to phosphoproteome signature that drive cardioprotection in synthetic torpor. Synthetic torpor induction by MPA neuronal activation induces changes to kinase signatures in cardiac tissue which are associated with cell survival/stress, cell cycle/death and cellular function. Activated kinases are shown in green and inhibited kinases in blue. Arrows indicate kinases that act downstream of the ones above. Abbreviations: CK2A2, casein kinase II alpha II subunit; AMPKA1, AMP-activated protein kinase alpha 1; MAPK10/JNK3, Mitogen-activated protein kinase 10/ c-Jun N terminal kinase; ACVR1, Activin A receptor type 1; MAPK12/13/14, Mitogen-activated protein kinase 12/13/14; MK2/3/5, Mitogen-activated protein kinase-activated protein kinase 2/3/5; AKT1, AKT serine/threonine kinase 1; p70S6K, 70 kDa ribosomal protein s6 kinase; CAMKIIA/G, calcium/calmodulin-dependent protein kinase II A/G; TP53, Tumour protein p53; PI3K, Phosphoinositide 3-kinase; mTOR, mechanistic target of rapamycin.

Several MAPK kinases were inhibited during synthetic torpor, including MAPK10/JNK3, MAPK12-14 and downstream MAPKAPK2 (MK2), MAPKAPK3 (MK3), & MAPKAPK5 (MK5). MAPK signalling is fundamental for cell survival, inflammation and regulating apoptosis. Cell stress, apoptosis and inflammation are strongly associated with ischemia-reperfusion injury and several studies have demonstrated pharmacological inhibition of p38 MAPK pathway reduced cardiac remodelling and infarct size post myocardial infarction injury^77^. Altered MAPKs have been reported in deep hibernation^78,79^, supporting the hypothesis that synthetic torpor in the rat is analogous to natural torpor/hibernation.

Tightly regulated calcium signaling within the heart is crucial for muscle contraction and maintaining a regular heart rhythm. Excessive CAMKII activation is linked to dysregulated calcium signaling, myocardial dysfunction and electrical instability^80^. Furthermore, in ischemia-reperfusion injury there is a disruption to calcium signaling during ischemia, followed by overload on reperfusion, leading to mitochondria dysfunction and cell death^81^. Here we demonstrate downregulated activity of CAMKIIA and CAMKIIG kinases, which is typically associated with improved contractility and reduced cell death in models of heart disease^80,82^. Changes to calcium handling and CAMKII signaling have also been observed in mammalian hibernators^83,84^. We also demonstrated a reduction in ACVR1 activity. ACVR1 is thought to mediate the development of cardiac hypertrophy and blocking this signaling pathway reduces pathological cardiac remodeling^85^.

These findings are also consistent with the GSEA reactome data (figure 4), with modulation of pathways associated with cell stress and survival (such as PI3/AKT/mTOR, MAPK and calcium signaling), and cell cycle and death (such as autophagy and apoptosis). We hypothesise that the activation of MPA neurons within the rat hypothalamus drives changes in a network of pathways within the cardiac tissue, that synergistically contribute to the cardioprotective effect of synthetic torpor (figure 6).

## Limitations

These experiments were conducted in an ex-vivo Langendorff model of ischaemia-reperfusion injury. While we have no reason to suppose that similar protection would not be observed in-vivo, we have not explicitly tested this. Synthetic torpor was applied prior to the ischaemia-reperfusion injury, it remains to be seen whether synthetic torpor applied after reperfusion can modulate the injury. Our phosphoproteme analysis was exploratory and hypothesis generating due to relatively small numbers of samples and the high demands of FDR-corrected statistics when analysing 1000s of phosphopeptides. The preoptic area of the hypothalamus is enriched with estrogen receptors^49^ and previous work by our group has demonstrated that the estrous cycle modulates fasting-induced torpor^86^. Our current work did not aim to investigate sex differences and is not powered to include this analysis. It should also be noted that the effect on oxygen consumption was not assessed in the thermoneutral synthetic torpor animals and we consequently cannot say whether a reduction in metabolic rate occurred in these animals.

Our study represents an important step towards harnessing torpor and synthetic torpor states for clinical translation. We have shown that species that do not naturally enter torpor can be induced to do so through targeted activation of neuronal populations that correspond to those responsible for natural torpor in the mouse. This state of synthetic torpor confers protection from ischaemia-reperfusion independent of body temperature or heart rate, and which persists in the isolated heart. It adds to a growing body of evidence that the central nervous system is able to induce protective adaptations in the heart^87–89^.

## Methods

### Rats

All studies had the approval of the local University of Bristol Animal Welfare and Ethical Review Board and were conducted in accordance with the UK animals (scientific procedures) act. Male and female Wistar rats (175-250 g) were obtained from Charles River and were housed at 21°C on a 12-hour light/dark cycle with ad lib access to food and standard rat chow (LabDiet).

### Viral vectors

AAV-CaMKIIa-hM3D(Gq)-mCherry was a gift from Bryan Roth (Addgene viral prep # 50476-AAV5) and was used at a titre of 1.7×10^13^ viral genome copies per ml. AAV-CaMKIIa-EGFP also a gift from Bryan Roth (Addgene viral prep # 50469-AAV5) and was used at a titre of 2.8×10^13^ viral genome copies per ml.

### Stereotaxic injection of viral vectors

Rats were anaesthetised with intraperitoneal injection of ketamine (5mg/100g, Vetalar; Pharmacia) and medetomidine (30μg/100g Domitor, Pfizer) until loss of paw withdrawal reflex. Additional intraperitoneal injections of anaesthetic were administered as needed to maintain surgical depth of anaesthesia. Core temperature was maintained using a servo-controlled heat pad and a rectal temperature probe (Harvard Apparatus). The planned incision site was shaved, the skin was cleaned with iodine solution, and sterile surgical technique was used throughout. Anesthetized rats were placed in a stereotaxic frame, the head was fixed in atraumatic ear bars, and skull position maintained by an incisor bar (David Kopf Instruments). Microcapillary pipettes were made from microcapillary glass (Sigma) on a vertical pipette puller (Harvard Apparatus). Pipettes were filled with mineral oil; then vector was back-filled using a robotic microinjector (Nano-W wireless capillary microinjector, Neurostar), producing a visible vector–mineral oil interface. The scalp was incised in the midline and burr holes made bilaterally with a drill attachment (Neurostar). Bilateral injections were made at depths of 8.5mm relative to the surface of the brain, +0.12mm from Bregma and 0.4mm from the midline. Each injection was 200nl and was delivered at a rate of 100nl/minute. The injection pipette remained in place for 5 minutes after the injection before slowly removing. Following vector injections, the wound was closed with nonabsorbable suture and dressed with antibacterial wound powder. Anaesthesia was reversed with atipamezole (1 mg/kg i.p, Antisedan, Zoetis), and meloxicam analgesia given subcutaneously (5mg/kg, Metacam, Boehringer Ingelheim). Rats were recovered in a heated chamber, then housed separately for three days following surgery and monitored daily.

### Implantation of telemeters

During the anaesthesia for stereotaxic CNS injections, a subset of rats was additionally implanted with telemetric temperature and electrocardiogram (ECG) probes into the abdomen (DSI ETA-F10 implant, Data Science International, Harvard Bioscience Inc). A vertical incision was made through the skin in the abdomen just below the xiphisternum approximately 2-3 cm in length and a smaller incision made to the skin at the right-hand side of the chest, for placement of the ECG electrode. Telemeter wires were passed under the skin and sewn into place using non-absorbable sutures. An intraperitoneal incision was made into the abdomen for placement of the telemetry implant, which was closed with absorbable sutures. Finally, the skin incisions were closed with non-absorbable sutures and dressed with antibacterial wound powder. Anaesthesia was reversed with atipamezole (1 mg/kg i.p, Antisedan, Zoetis), and meloxicam analgesia given subcutaneously (5mg/kg, Metacam, Boehringer Ingelheim). Rats were recovered in a heated chamber, then housed separately for three days following surgery and monitored daily.

### Induction of synthetic torpor

Intraperitoneal injection of clozapine-N-oxide dihydrochloride (CNO), a synthetic ligand that activates the DREADD hM3D_q_ (2 mg/kg, Tocris Bioscience, #6329), was used to induce synthetic torpor. For those rats that were not implanted with telemeters, synthetic torpor induction was confirmed using thermal imaging with a Flir C2 camera. Synthetic torpor was defined by a drop in core temperature below 35°C in telemetered animals, or surface temperature below 30°C in thermally imaged animals following CNO injection. Some MPA HM3DGq viral injections failed and animals did not enter synthetic torpor, the success rate for inducing synthetic torpor was 80%, so consequently 20% of the animals were excluded from the study.

### Telemetric monitoring of core body temperature, ECG and motor activity

Telemetry implants were switched on and connected to a receiver platform (DSI PhysioTel^TM^ receiver RPC-1, Data Science International, Harvard Bioscience Inc). Continuous telemetry data, including core body temperature, ECG and motor activity were acquired and recorded wirelessly using DSI Matrix 2.0 (MX2, Data Science International, Harvard Bioscience Inc) and Ponemah software (Ponemah v6.00, Data Science International, Harvard Bioscience Inc). Recordings were made from animals in their home cages for data at room temperature. For the thermoneutral condition maintaining control rats in an ambient temperature of 30°C caused them to become hyperthermic, hence 28°C was used for these animals, while MPA^Gq^ rats remained euthermic at 30°C following CNO injection. Post recording, ECG, temperature and activity data were filtered to remove noise and analysed in Ponemah software. The R wave component of the ECG recordings was detected and used to determine the heart rate in bpm.

For some animals, surface body temperature in place of implanted telemetry was used to confirm entry into synthetic torpor. To do this, rats were moved into a custom built Perspex 32 × 42 × 56 cm cage and surface body temperature was recorded using an infrared thermal imaging camera placed above the cage (Flir C2, www.flir.co.uk).

### Measurement of oxygen consumption

Indirect calorimetry was performed using a CaloBox (Phenosys)^90^, with a gas flow rate of 75 litres per hour and a sampling rate of 0.25Hz. Rats were first acclimated to the recording chamber for 30 minutes over 3-5 days before undergoing CNO injection. Data was discarded for 10 minutes following CNO injection to allow the CaloBox to re-equilibriate. Food and water were available throughout the recording session.

### Ex vivo model of cardiac ischemia-reperfusion injury

90 min post CNO administration, an intraperitoneal injection of heparin was delivered (1,000 IU, Sigma-Aldrich, #H3393). Animals were deeply anesthetized with isoflurane (5%) which was delivered in a constant flow of oxygen (2 l/min). Following loss of pedal reflex, animals were sacrificed by cervical dislocation. A thoracotomy was performed and beating hearts were removed by sectioning the aorta. Hearts were quickly placed into ice-cold Krebs-Henseleit Buffer (KHB) (containing 120 mM NaCl, 25 mM NaHCO_3_, 11 mM Glucose, 1.2 mM KH_2_PO_4_, 1.2 mM MgSO_4_, 4.8 mM KCl and 1.2 mM CaCl_2_). The aorta was then cannulated (secured into place using suture threads) and hearts were perfused by the Langendorff method (Langendorff, 1898) on an isolated heart system apparatus. Hearts were perfused with carbonated KHB, maintained at 37°C, at a rate of 10 ml/min. ECG was continuously recorded by Ag electrodes placed near the apex of the heart. After a 30-minute stabilization phase, the hearts were then subjected to 30 minutes of global ischemia, whereby perfusion was stopped, and hearts were submerged into a water-jacketed bath of KHB held at 37°C. After ischemia, the hearts were reperfused for 60 minutes (figure 2A). Exclusion criteria for experiments included time between excision of the heart and start of ex vivo perfusion with KHB of more than 2 minutes, more than 2 aortic cannulation attempts, and heart rate under 200 bpm at the end of the equilibration period. A total of 8 animals were excluded based on these criteria.

### Heart infarction assessment

Following 60 minutes of reperfusion, the hearts were perfused with 2.3.5-Triphenyltetrazolium Chloride (TTC) (Phytotech, USA) to demarcate infarcted tissue (unstained by TTC) and viable tissue (stained by TTC)^91^. 1% TTC solution at 37°C was perfused through the aortic cannula at a rate of 2 ml/min and then hearts were submerged in 1% TTC for a further 15 mins at 37°C, before being fixed in 4% paraformaldehyde for 24 hours at 4°C. Hearts were then transversally sliced into 3 mm thick sections and each side of the sections were imaged using a flatbed scanner. ImageJ (NIH, USA) was used to outline and calculate the infarcted area and whole heart area (excluding ventricular chambers). Both infarcted and viable areas were averaged across each side for each heart section and infarct area was expressed as a percentage dividing total infarcted area by whole heart area. The assessor was blinded to the animal group.

### Mapping of virally transfected MPA hypothalamic neurons

#### Immunofluorescence

Brains were extracted and post-fixed for 48 hours with 4% PFA at 4°C and then incubated in PBS with 30% sucrose and 0.03% sodium azide for a further 48 hours. Brains were embedded in tissue freezing medium and sectioned on a freezing microtome into 40 µm coronal sections and adhered to SuperFrost Plus slides (Epredia). A hydrophobic barrier was drawn around each slide using an ImmEdge Pen (Vector Laboratories) and allowed to dry at room temperature for 15 min. Sections were washed 3 times for 5 min in PBS and then blocked with 0.25% Bovine Serum Albumin (BSA) and 10% Normal Donkey Serum (NDS) in 0.3% Triton-X PBS (PBS-T) for 4 hours at room temperature. They were then incubated with primary antibody (anti-mCherry, 1:2000 dilution, BioVision, #5993) diluted in 0.25% BSA and 2% NDS in PBS-T overnight at room temperature. Slides were then washed 3 times for 5 min in PBS, followed by incubation with the secondary antibody (Alexa Fluor 594 donkey anti-rabbit, 1:1000 dilution, Abcam, #A-21207) diluted in 0.25% BSA in PBS-T for 4 hours at room temperature. Finally, slides were washed 3 times for 5 min in PBS and mounted with Fluoroshield with DAPI (Sigma Aldrich).

#### Epifluorescence imaging & mapping

Sections were imaged using a Leica DM4000B widefield microscope with a 20x magnification 0.5 numerical aperture objective, with a 515-561nm excitation filter, dichroic mirror 580nm, and a 590nm emission filter. The anatomical extent of virally transduced cells was mapped independently by two individuals and agreed by consensus using the Allen brain atlas as reference.

### Characterisation of neurons driving this torpor-like state

#### Fluorescence in situ hybridization (ISH)

Brains were extracted and immediately frozen in 2-methylbutane and stored at -80°C until use. Brains were embedded in tissue freezing medium (Cryomatrix^TM^, ThermoFisher) and sliced on a cryostat into 20µm coronal sections, adhered to SuperFrost Plus slides (Epredia) and stored at -20°C overnight before undergoing the standard RNAscope Fluorescent Multiplex Assay for fresh frozen tissue (ACD Bio, Biotechne).

For the RNAscope ISH, mounted sections were fixed in 4% PFA for 15 min at room temperature, then dehydrated in increasing concentrations of ethanol (50%, 75%, then 100%) for 5 minutes in each. Sections were left overnight in a 2^nd^ application of 100% ethanol. 2 sections per animal were selected for processing based on mCherry expression within the MPA and underwent 1 of 2 protocols, protocol 1: markers of excitatory and inhibitory neurons, and Protocol 2: markers that have been associated with neurons controlling mouse torpor (see table 1 for probes used). Dehydrated sections were air-dried for 10 min at room temperature, treated with hydrogen peroxide for 10 minutes at room temperature, washed with distilled water and then treated with protease IV for 12 min. The RNAscope probes were then applied to each slide and left to hybridize for 2 h at 40°C in a humidified chamber. Following hybridization, slides were washed with wash buffer twice for 2 minutes at room temperature. AMP1 was applied to each slide and left to incubate in a humidified oven for 30 minutes at 40°C, then washed again twice for 2 minutes. This amplification process was repeated for AMP2 and AMP3. Next, slides were washed twice with wash buffer for 2 minutes at room temperature and HRP-C1 was applied to sections for 15 minutes at 40°C in a humidified chamber. Slides were rinsed again with wash buffer for 2 minutes at room temperature and an Opal dye was incubated on the slides for 30 minutes at 40°C in a humidified chamber. Following this, slides were washed with wash buffer for 2 minutes at room temperature and the HRP blocker was applied to sections for 15 minutes at 40°C in a humidified chamber. This process was repeated for HRP-C2, C3 and C4 with different Opal dyes for each. Finally, the sections were rinsed in wash buffer for 2 minutes at room temperature and coverslipped with fluoroshield with DAPI and left to dry at room temperature overnight, before being stored at 4°C.

**Table 1:**
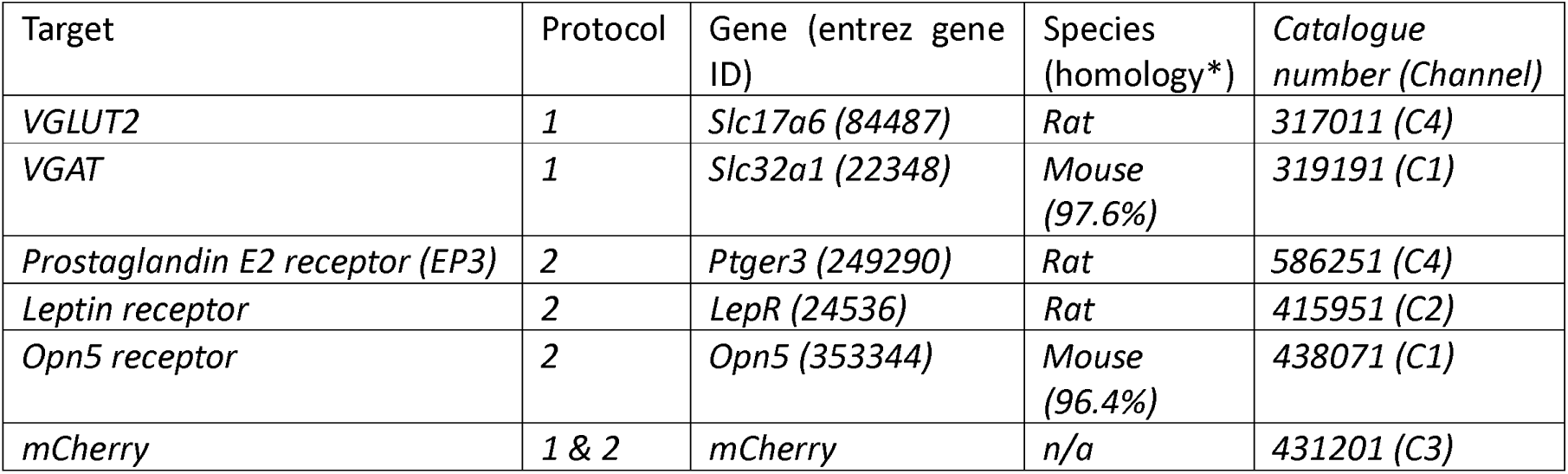
RNAscope probes used for ISH. *https://rddc.tsinghua-gd.org/.

| Target | Protocol | Gene (entrez gene ID) | Species (homology*) | Catalogue number (Channel) |
| --- | --- | --- | --- | --- |
| VGLUT2 | 1 | Slc17a6 (84487) | Rat | 317011 (C4) |
| VGAT | 1 | Slc32a1 (22348) | Mouse (97.6%) | 319191 (C1) |
| Prostaglandin E2 receptor (EP3) | 2 | Ptger3 (249290) | Rat | 586251 (C4) |
| Leptin receptor | 2 | LepR (24536) | Rat | 415951 (C2) |
| Opn5 receptor | 2 | Opn5 (353344) | Mouse (96.4%) | 438071 (C1) |
| mCherry | 1 & 2 | mCherry | n/a | 431201 (C3) |

#### Confocal imaging

Brains were processed as indicated in Fluorescence in situ hybridization section and imaged within 2 weeks of processing. Sections containing MPA were imaged on a Leica SP8 AOBS confocal laser scanning microscope attached to a Leica DMi8 inverted epifluorescence microscope using HyD detectors (“hybrid” SMD GaAsP detectors) with 405 nm diode and white light lasers. A 40x oil immersion lens with NA of 1.3 (HC PL APO CS2 lens). Tiled MPA images were taken and stitched using LASX software (Leica) and channels were imaged sequentially to avoid optical crosstalk.

#### In situ hybridization image analysis

Images were z projection compressed in ImageJ. A 50-pixel radius rolling ball background subtraction was applied to each image. An automated image processing pipeline was used to count DAPI-and RNA-positive nuclei. An Otsu threshold was applied to the DAPI image and overlapping nuclei were separated using the watershed method. Cell nuclei were then filtered by size (20-200µm^2^) and counted and ROIs for each cell nuclei were saved in ImageJ. ROIs and cell identities created from the DAPI image were then overlayed on the ISH images and cell nuclei positive for RNA were counted. A minimum of 200 cells from each brain section were manually assessed to determine a threshold for nuclei positive for the different RNA transcripts and these thresholds were then applied to the remaining cells.

### Proteomics

#### Tissue preparation

90 minutes post CNO administration, animals were deeply anesthetized with isoflurane (5%) delivered in a constant flow of oxygen (2 l/min) and sacrificed by cervical dislocation following loss of pedal reflex. A thoracotomy was performed and beating hearts were removed by sectioning the aorta. Hearts were quickly placed into ice-cold cardioplegia buffer (7.4 pH, containing 20 mM HEPES, 15 mM Glucose, 15 mM KCl, 7.8 mM MgSO_4_ and 1.2 mM CaCl_2_) and agitated to remove blood. Hearts were then immersed and immediately frozen in 2-methylbutane in liquid nitrogen and stored at - 80°C until use. For protein sample preparation, hearts were thawed gently on ice and 50-100 mg sample was cut from the apex of each heart and placed into 10 µL of RIPA buffer and homogenized (3 × 15 second bursts at 10,000 RPM). Samples were then centrifuged at 10,000 g for 10 minutes at 4°C. Supernatant was transferred into a new tube and protein concentration was determined with the BCA Protein Assay kit. Protein samples were diluted to a final concentration of 2 mg/mL in RIPA buffer and stored at -80°C.

#### TMT labelling, high pH reverse-phase chromatography and phosphor-peptide enrichment

Protein samples were analysed using Mass Spectrometry (MS) using tandem mass tag (TMT) (Proteomics facility at University of Bristol). Aliquots of 100 µg of each sample were digested with 2.5 µg trypsin at 37°C overnight and labelled with TMT (TMTpro) sixteen-plex reagents and the labelled samples were then pooled.

For total proteome analysis, an aliquot of 100 µg of the pooled sample was desalted using a SepPak cartridge (Waters). Eluate from the SepPak cartridge was evaporated and resuspended in buffer A (20 mM ammonium hydroxide, pH 10) prior to fractionation by high pH reversed-phase chromatography using an Ultimate 3000 liquid chromatography system (Thermo Fisher Scientific). Briefly, the sample was loaded onto an XBridge BEH C18 Column (130Å, 3.5 µm, 2.1 mm X 150 mm, Waters, UK) in buffer A and peptides eluted with an increasing gradient of buffer B (20 mM Ammonium Hydroxide in acetonitrile, pH 10) from 0-95% over 60 minutes. The resulting fractions (concatenated into 20 in total) were evaporated to dryness and resuspended in 1% formic acid prior to analysis by nano-LC MS/MS using an Orbitrap Fusion Lumos mass spectrometer (Thermo Scientific).

For phosphoproteome analysis, the remainder of the TMT labelled pooled sample was also desalted using a SepPak cartridge and eluate evaporated and then subjected to TiO2-enrichement according to manufacturer’s protocol. The flow-through and washes from the TiO2-based enrichment were then subjected to FeNTA-based phosphopeptide enrichment according to the manufacturer’s instructions (Pierce). The phospho-enriched samples were again evaporated to dryness and then resuspended in 1% formic acid prior to analysis by nano-LC MS/MS.

#### Nano-LC Mass Spectrometry

High pH RP fractions (Total proteome analysis) or the phospho-enriched fractions (Phospho-proteome analysis) were further fractionated using an Ultimate 3000 nano-LC system in line with an Orbitrap Fusion Lumos mass spectrometer.

Peptides in 1% (vol/vol) formic acid were injected onto an Acclaim PepMap C18 nano-trap column (Thermo Scientific). After washing with 0.5% (vol/vol) acetonitrile 0.1% (vol/vol) formic acid peptides were resolved on a 500 mm × 75 μm Acclaim PepMap C18 reverse phase analytical column (Thermo Scientific) over a 150 min organic gradient, using 7 gradient segments in solvent B (aqueous 80% acetonitrile in 0.1% formic acid). (Gradients: 1-6% over 1 min, 6-15% over 58 min, 15-32% over 58 min, 32-40% over 5 min, 40-90% over 1 min, held at 90% for 6 min and then reduced to 1% over 1 min) with a flow rate of 300 nl/min. Peptides were ionized by nano-electrospray ionization at 2.0kV using a stainless-steel emitter with an internal diameter of 30 μm (Thermo Scientific) and a capillary temperature of 300°C.

All spectra were acquired using an Orbitrap Fusion Lumos mass spectrometer controlled by Xcalibur 3.0 software and operated in data-dependent acquisition mode using an SPS-MS3 workflow. FTMS1 spectra were collected at a resolution of 120,000, with an automatic gain control (AGC) target of 200 000 and a max injection time of 50 ms. Precursors were filtered with an intensity threshold of 5000, according to charge state (to include charge states 2-7) and with monoisotopic peak determination set to Peptide. Previously interrogated precursors were excluded using a dynamic window (60 s +/-10 ppm). The MS2 precursors were isolated with a quadrupole isolation window of 0.7 m/z. ITMS2 spectra were collected with an AGC target of 10,000, max injection time of 70 ms and CID collision energy of 35%.

For FTMS3 analysis, the Orbitrap was operated at 50,000 resolution with an AGC target of 50,000 and a max injection time of 105 ms. Precursors were fragmented by high energy collision dissociation (HCD) at a normalised collision energy of 60% to ensure maximal TMT reporter ion yield. Synchronous Precursor Selection (SPS) was enabled to include up to 10 MS2 fragment ions in the FTMS3 scan.

#### Data processing

The raw data files were processed and quantified using Proteome Discoverer software v2.4 (Thermo Scientific) and searched against the UniProt Rat database (downloaded July 2025: 47902 entries) using the SEQUEST HT algorithm. Peptide precursor mass tolerance was set at 10 ppm, and MS/MS tolerance was set at 0.6 Da. Search criteria included oxidation of methionine (+15.995 Da), acetylation of the protein N-terminus (+42.011 Da), methionine loss from the protein N-terminus (−131.04 Da) and methionine loss plus acetylation of the protein N-terminus (−89.03 Da) as variable modifications and carbamidomethylation of cysteine (+57.0214) and the addition of the TMTpro mass tag (+304.207) to peptide N-termini and lysine as fixed modifications. For the Phospho-proteome analysis, phosphorylation of serine, threonine and tyrosine (+79.966) was also included as a variable modification. Searches were performed with full tryptic digestion and a maximum of 2 missed cleavages were allowed. To ensure a high confidence in the identified phosphopeptides to the matched proteins, the reverse database search option was enabled and all data was filtered to satisfy false discovery rate (FDR) of 5%.

#### Phosphoproteome analysis

Phosphoproteome data was processed from a single dataset containing log2-adjusted, normalised phosphopeptide abundances. Phosphosites were retained if a residue position could be unambiguously parsed and, where available, localisation probability was ≥75%. Intensities were collapsed to one value per phosphosite by averaging across peptides mapping to the same site. Site-level quality control was performed using multiple orthogonal metrics, including missingness, inter-sample correlation, PCA distance, and distributional properties. These metrics were combined into an objective composite score used for diagnostic purposes only; no samples were excluded a priori.

Differential phosphorylation was assessed using limma^41^ with empirical Bayes moderation. Initial heatmap and PCA analysis of the most differentially enriched phospho-sites revealed a clear batch effect (likely due to one batch undergoing a freeze-thaw cycle prior to protein isolation, supplementary figure 4). Limma modelling^41^using batch as a covariate effectively controlled the effect, with samples clustering by experiment group (figure 4). Genes were ranked for GSEA using the most extreme site-level moderated t-statistic per gene from a limma model including batch as a covariate. This ranking incorporates both effect size and variance and avoids over-weighting noisy phosphosites. Gene-level pathway enrichment was performed using Reactome PA^42,92^, ranking genes by site-derived t statistic. Phosphosite-level kinase and pathway enrichment was performed using directional PTM-SEA implemented via fgsea^45^. Gene identifier mapping was performed using biomaRt^43^ and AnnotationDbi^44^. Cross-species phosphosite mapping was performed using pairwise protein sequence alignment implemented in Biostrings using pwalign^93,94^. Rat phosphosites were mapped to human orthologues via gene orthology and full-length protein sequence alignment, retaining only sites with matched residues and position. Directional PTM-SEA was then performed using fgsea^45^ with site-level t-statistics as the ranking metric and PTMsigDB kinase signatures. Kinase enrichment results were visualised using volcano plots, dotplots, and heatmaps. All enrichment thresholds were treated as hypothesis-generating and are reported as such.

## Statistical analysis

Descriptive statistics including mean ± standard deviations are reported in the main text. Statistical details for experiments can be found in figure legends, including n numbers, statistical test used and level of significance. Data distributions were visually inspected and tested for normality using Shapiro-Wilk test. All data were normally distributed. One-way analysis of variance (ANOVA) was used for multiple group comparisons, followed by Sidak’s or Holm-Sidak’s post hoc test to identify significant differences as reported in the text and figure legends. There were 2 independent factors, viral vector construct (MPA^EGFP^ and MPA^Gq^) and ambient environment temperature (room (21-22°C) or thermoneutral (30°C)). The dependent factors were core body temperature, heart rate, activity and heart infarct size. Activity data was analysed using a repeated measures ANOVA. A two-tailed paired t-test was used to analyse the oxygen consumption data, to assess within animal changes, pre-and post-CNO injection. All analyses were performed in GraphPad Prism (Version 10.4.1).

## Data Availability

Phosphoproteome data alongside analysis scripts were deposited here: https://doi.org/10.5061/dryad.v6wwpzh9r

## Acknowledgements

We would like to thank undergraduate students Aditya Sethi and Hiyaan Rizvi for their help with processing the proteomics samples and Zeynab Mohamed for her help with collecting some of the mCherry immunohistochemistry data.

## Supplementary figures

**Supplementary figure 1.**
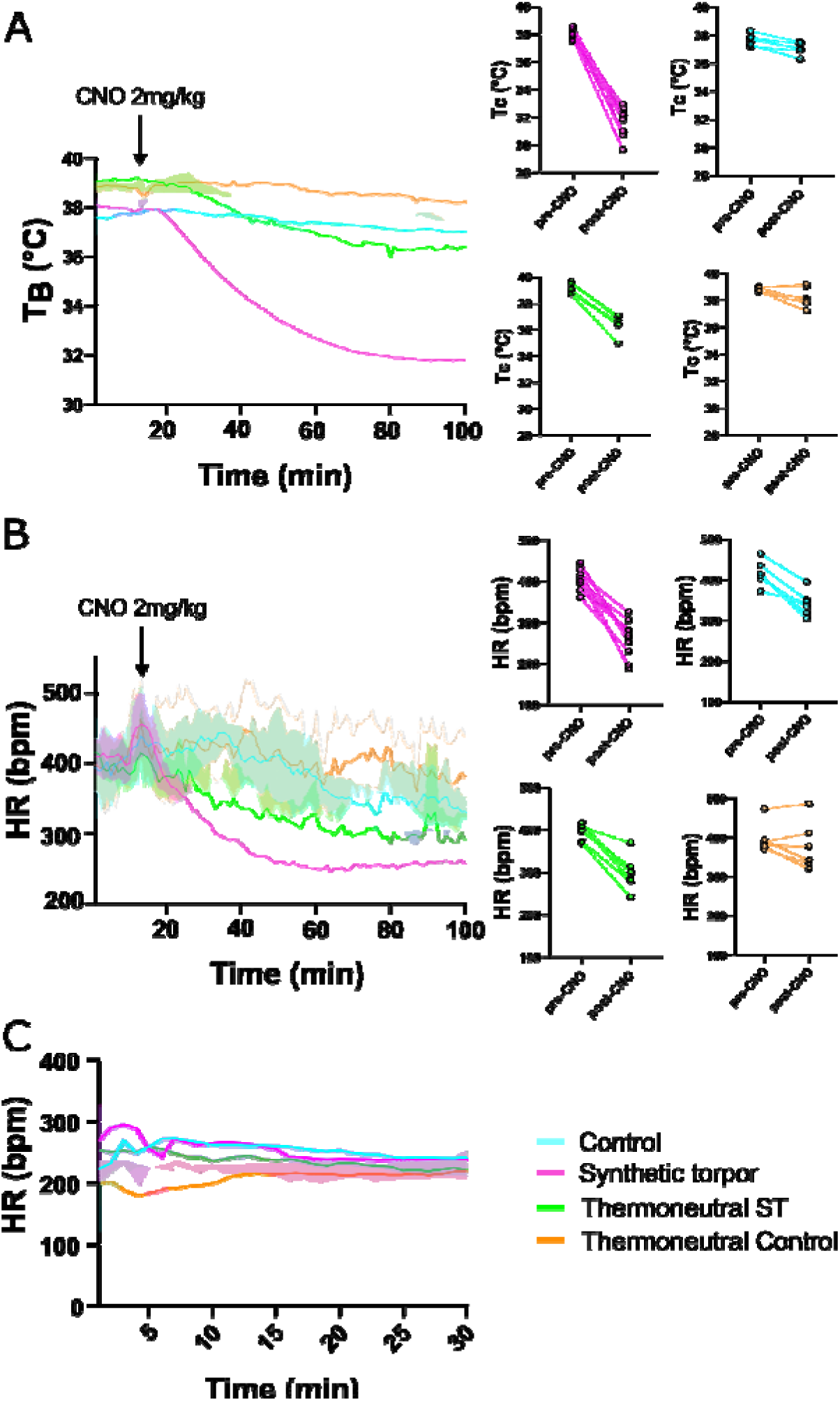
Pre-Langendorff *in vivo* physiological changes (A,B) and *ex vivo* heart rates (C). Core temperature (A) and heart rate (B) *in vivo* prior to excision of heart for Langendorff preparation. A,B, n = 6 controls, 10 synthetic torpor, 6 thermoneutral synthetic torpor, & 6 thermoneutral controls. Pre (10-20 min) and post (90-100 min) CNO averages for T_b_ and HR in control, torpor, thermoneutral synthetic torpor and thermoneutral controls. (C) No significant differences in *ex vivo* HR were observed between the four groups during the 30-minute equilibration period of the Langendorff assay (F(3,19) = 1.35, *p* = 0.29). Controls (n = 16), torpor (n = 12), thermoneutral synthetic torpor (n = 5), & thermoneutral controls (n = 6). Data shows mean and 95% confidence interval. Abbreviations: ST, synthetic torpor; T_b_, core body temperature; HR, heart rate; bpm, beats per minute.

**Supplementary figure 2.**
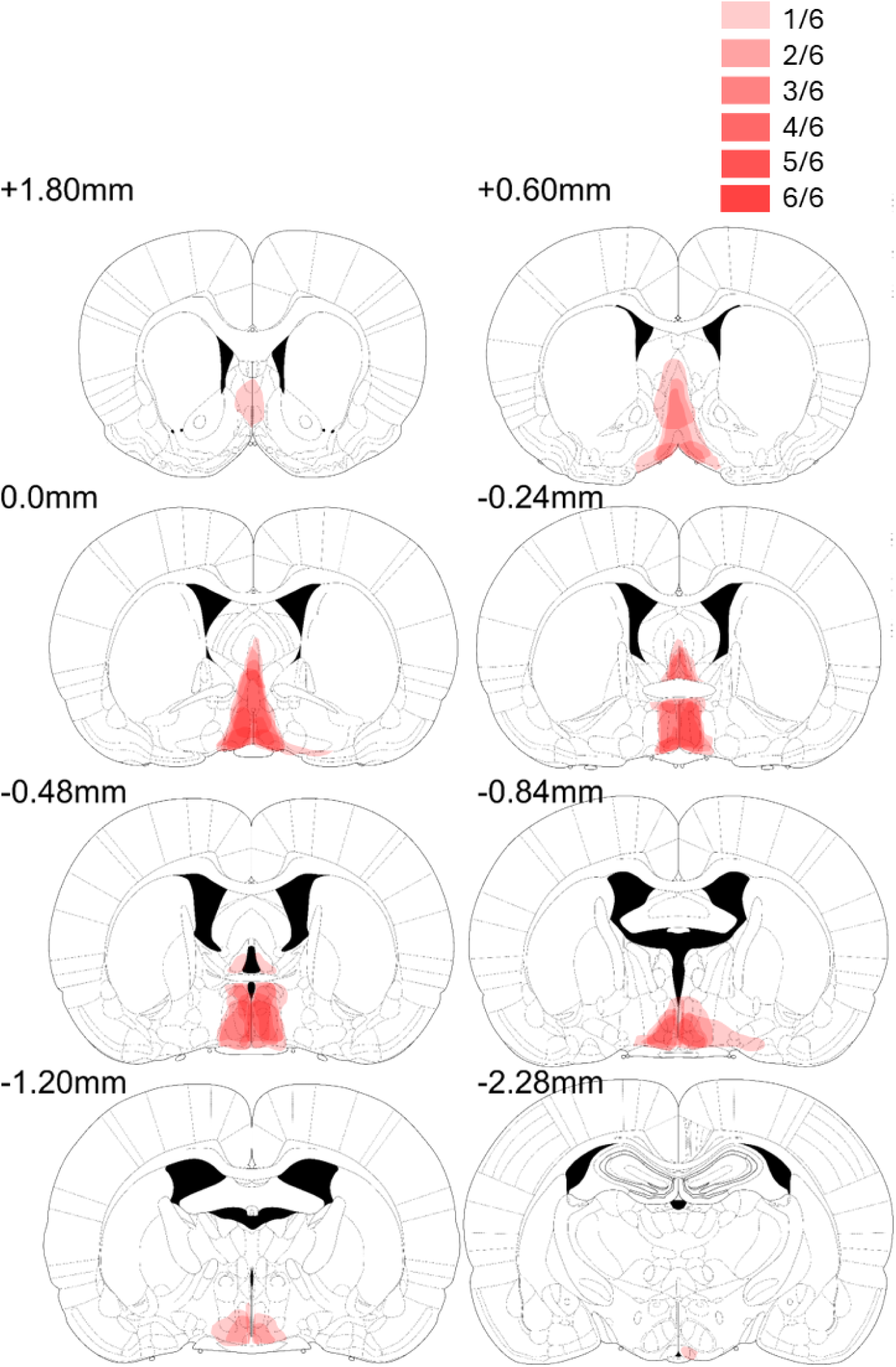
mCherry viral vector mapping demonstrating the transduction throughout the rat brain. Mapping shows viral vector transduction in 6 animals from bregma +1.80 to -2.28 mm.

**Supplementary figure 3.**
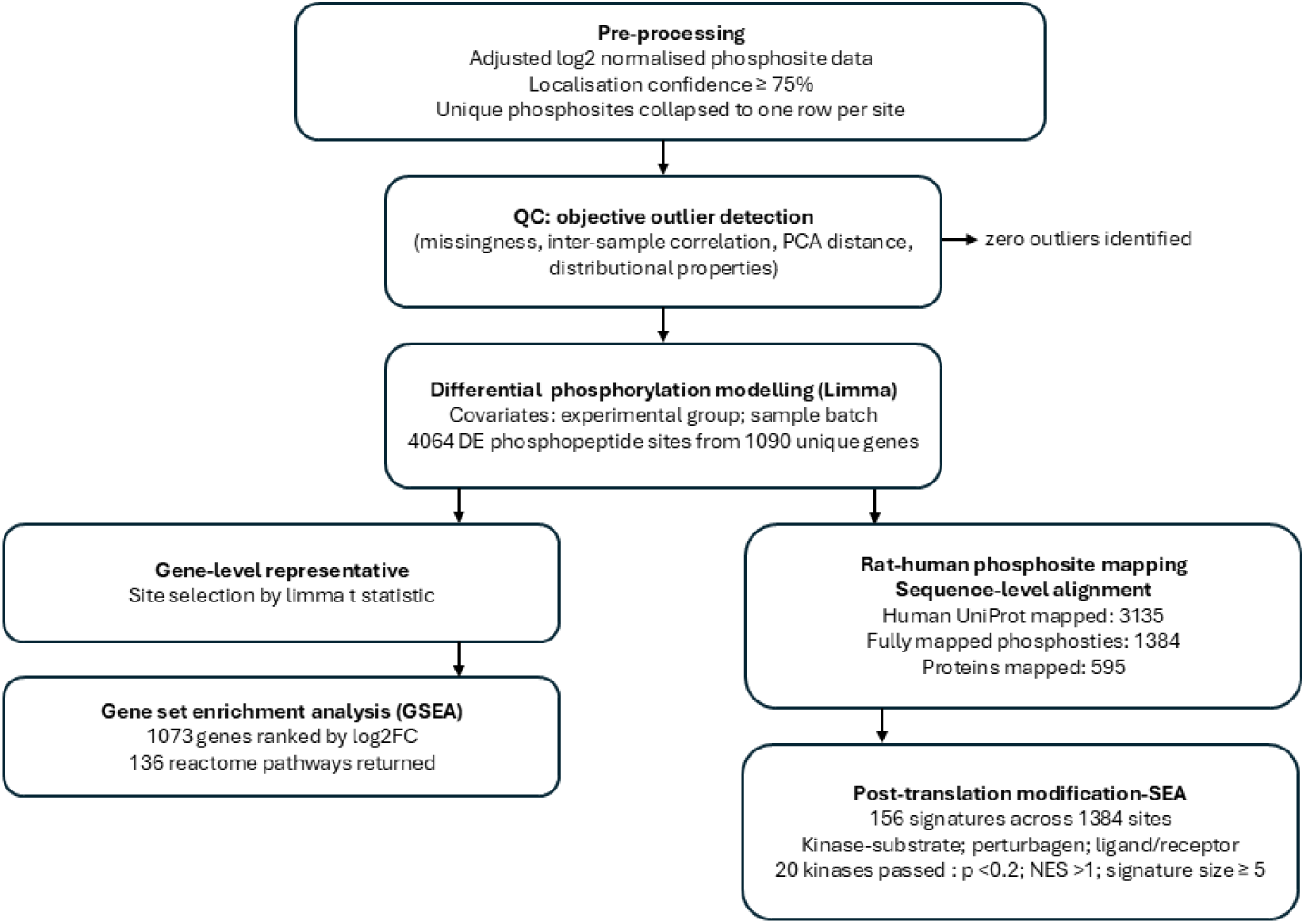
Phosphoproteomics workflow

**Supplementary figure 4.**
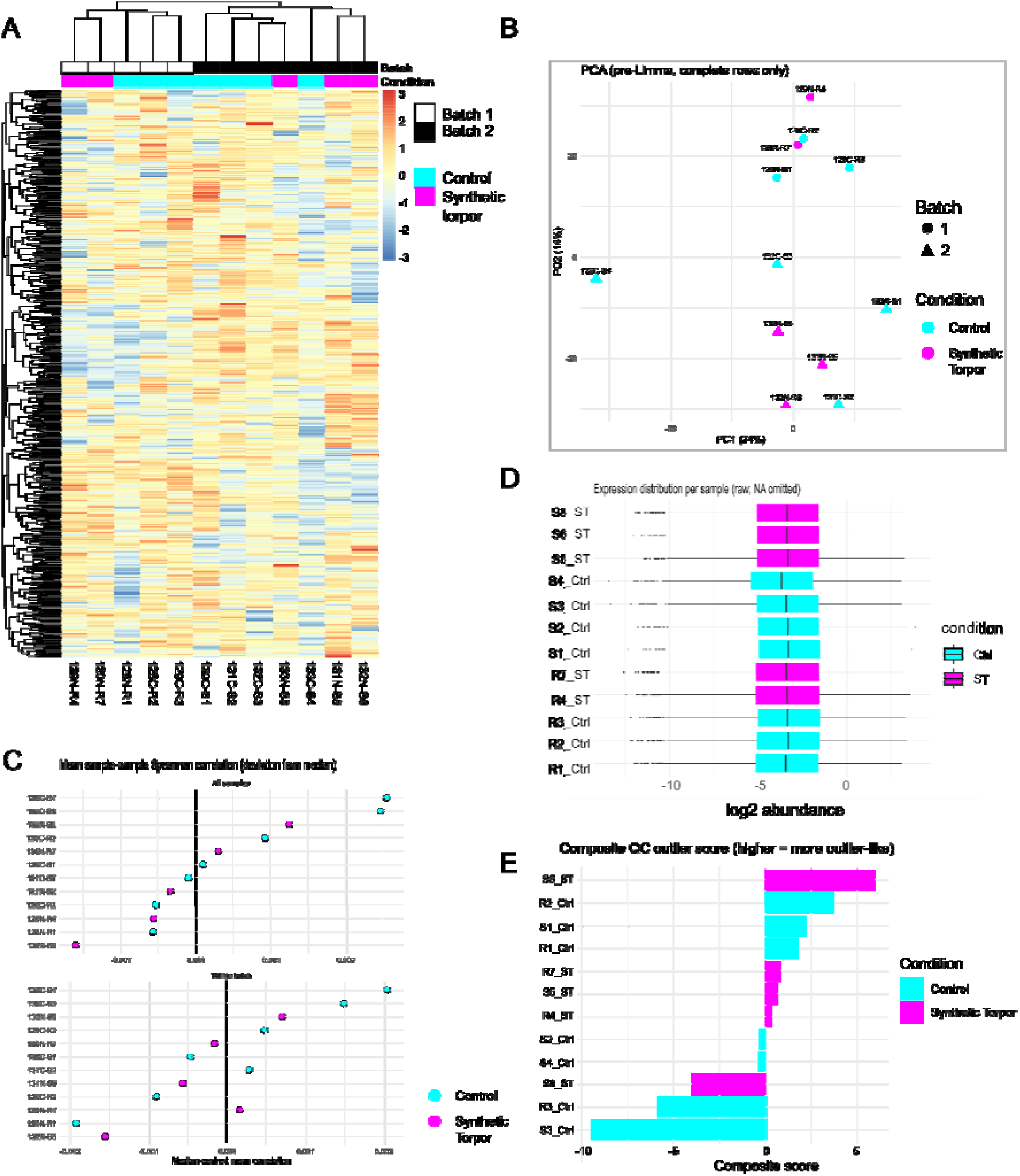
proteomics QC metrics. Heat map of 600 most variable phosphosites prior to limma modelling and batch correction (A) and principal component analysis of the same uncorrected log2 normalised phosphosites (B) both showing clear batch effect. High sample to sample correlation (all samples, top panel and within batch, bottom panel), indicating subtle biological effects and no outlier samples or process-specific effects (C). Sample-wise expression distributions assessed by computing the median and interquartile range (IQR) of log-transformed phosphoprotein intensities for each sample confirm no global shifts in signal intensity or variability prior to downstream modelling (D). Composite QC outlier scores confirms no outliers (E).

**Supplementary figure 5.**
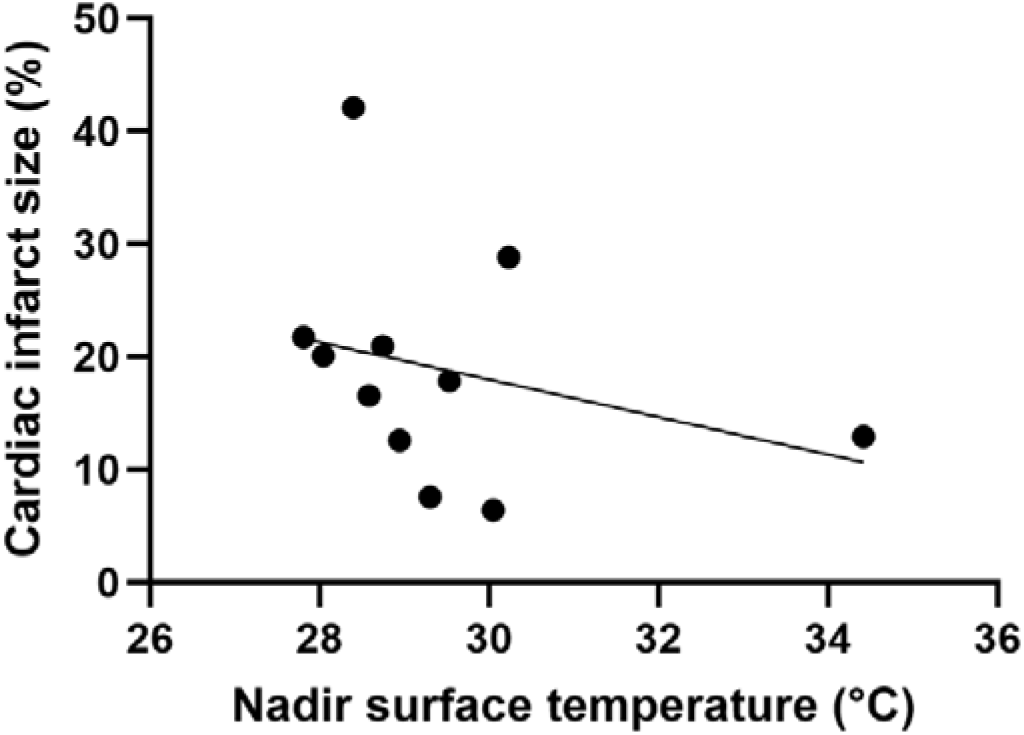
Relationship between nadir surface body temperature during synthetic torpor and cardiac infarct size. Linear regression revealed no significant relationship between the nadir surface temperature during synthetic torpor and cardiac infarct size *F*(1,9) = 0.885, *p* = 0.3714, *R*^2^ *=* 0.089.

